# Improved Monoisotopic Mass Estimation for Deeper Proteome Coverage

**DOI:** 10.1101/2020.06.03.131003

**Authors:** Ramin Rad, Jiaming Li, Julian Mintseris, Jeremy O’Connell, Steven P. Gygi, Devin K Schweppe

## Abstract

Accurate assignment of monoisotopic peaks is essential for the identification of peptides in bottom-up proteomics. Misassignment or inaccurate attribution of peptidic ions leads to lower sensitivity and fewer total peptide identifications. In the present work we present a performant, open-source, cross-platform algorithm, Monocle, for the rapid reassignment of instrument assigned precursor peaks to monoisotopic peptide assignments. We demonstrate that the present algorithm can be integrated into many common proteomics pipelines and provides rapid conversion from multiple data source types. Finally, we show that our monoisotopic peak assignment results in up to a two-fold increase in total peptide identifications compared to analyses lacking monoisotopic correction and a 44% improvement over previous monoisotopic peak correction algorithms.

## Introduction

Correct assignment of precursor masses is essential for robust determination of peptide spectral matches. The effects of misassigned precursor monoisotopic masses can severely degrade the identification rates of proteomic samples. With the increased speed of acquisition for modern instruments, low abundance and coeluting peptide masses can be missassigned prior to analysis. This error may be slight ppm shifts or large isotope shifts.

Monoisotopic peak estimation and correction has been a challenge for peptide spectral matching for decades^1, 2^. Several groups have developed methods to correct for monoisotopic peak assignment mistakes before^2-6^. Notably RawConverter, a .NET converter for Thermo Raw files can re-assign monoisotopic peaks using a linear optimization followed by chi-squared distance and cosine similarity to recalculate peak envelopes^2^. Based on total peptide identifications, RawConverter was reported to outperform previous monoisotopic peak correction algorithms by 14-46%. While monoisotopic peak re-assignment did dramatically improve the total peptide and protein identifications, we note that the success rate (total peptide count divided by the total MS2 spectra) for data dependent analyses (DDA) was often less than 40% even with RawConverter. We therefore set out to determine if further optimizing the monoisotopic peak assignment for DDA could improve success rates and thereby capture more peptides and proteins from a given set of spectra.

To attempt to address the low success rates we developed a new, cross-platform C# program termed Monocle. Monocle enabled both file conversion to common spectral formats (mzML^7^, mzXML^8^) and monoisotopic peak correction for improved detection sensitivity. Monocle has been released as a cross-platform, open-source project for integration into existing proteomics pipelines with both a core method library as well as a command line interface and graphical user interface.

We tested the utility of Monocle to improve the sensitivity of peptide spectral matching and protein identification of yeast and human peptides either as label-free or TMT-labeled proteomes^9^. Monocle peak correction increased valid peptide spectral matches – those passing a 1% peptide and protein FDR filter – up to 97% compared to raw data (no correction) and up to a 44% improvement over RawConverter. We go on to show that Monocle’s monoisotopic peak correction improved sensitivity for modern acquisition methods, e.g. those that use advanced precursor detection or ion mobility separations.

## Methods

### Sample Preparation

Human (HeLa) and yeast (BY4742) cell pellets were lysed (8 M urea, 50 mM EPPS pH 8.5, 150 mM NaCl, Roche protease inhibitor tablet) by syringe lysis and bead beating, respectively. Lysates were cleared via centrifugation, and the protein component was isolated by chloroform-methanol precipitation. Proteins were digested with LysC (Wako) overnight at room temperature followed by a 6-hour digestion with trypsin (Promega) at 37°C. An aliquot of digested HeLa peptides was subsequently labelled with TMT reagents at a 1:1 ratio across 10 channels. Labelled and label-free peptides were desalted using a C_18_ SepPak cartridge (Waters) prior to LC-MS/MS analysis. Samples ready for analysis were stored at - 80°C.

### Mass Spectrometric Data Acquisition and Analysis

Clean peptides were resuspended in 5% ACN/2% formic acid prior to loading on a 35cm, in-house pulled C18 column (100 μm ID, Thermo Accucore, 2.6 μm, 150 Å). Peptides were eluted and injected into a Thermo Fusion Lumos instrument and analyzed with a Top10 MS2 method using the low-resolution ion trap (FT-MS1: 120,000 R, 50 ms max injection time, AGC target of 1e5; IT-MS2 scans: rapid scan speed, 50 ms max injection time, AGC target of 2e4, normalized CID collision energy of 35% with 10 ms activation time, and 0.5 Th isolation width; HR-MS2 scans: 15,000 R, 50 ms max injection time, AGC target of 1e5, normalized HCD collision energy of 28 or 35, and 0.5 Th isolation width), unless otherwise noted. For FAIMS analyses, the compensation voltage was cycled through CV=-40/-60/-80V with a dispersion voltage of -5000V and all electrode temperatures set to 100°C.

Raw spectra were converted to mzXML via either Monocle, RawConverter, or an in-house, RawFileReader mzXML converter^2, 8^. Spectra were searched against Uniprot databases for yeast (Uniprot, 03-24-2020) or human (Uniprot, 02-25-2020) using the Comet search algorithm with default parameters except for: precursor mass tolerance of 50ppm, protease used was Trypsin/P^10^. For TMT-labelled samples searches were performed with the following modifications accounted for: variable Met oxidation (+15.99491), static Cys carboxyamido-methylation (+57.02146), and static TMT on Lys and peptide N-termini (+229.16293). Peptide spectral matches were filtered to a peptide and protein FDR less than 1% (valid PSMs)^11, 12^. Data were analyzed using R 3.6.3packages *ggplot2*^*13*^ and *dplyr*^*14*^. Linear regression was performed with R’s *lm* function (*stats* package). The matrix of sample analyses can be found in Table S1. Unless otherwise noted, values are presented with their standard deviation for replicate runs of the same sample type.

### Development and Open-source Implementation

Monocle was developed as a single C# dynamic link library (.Net Core 3.1). This library has been integrated into a command line interface (CLI) and a graphical user interface (GUI) projects. All three projects are available as an open source repository under the GPL-3.0 license (https://opensource.org/licenses/GPL-3.0). The source code can be found here: https://github.com/gygilab/Monocle. The use of the .NET Core framework ensures that Monocle can be built in either Windows or Linux environments for deployment in a wide variety of proteomics pipelines.

### Spectral Input and Pre-processing

Monocle can process common spectral file formats as input, including: Thermo RAW files, mzXML and mzML. Input files are read in either using custom code to read markup language files or the .Net Core build of RawFileReader to read RAW files. Spectra are read into a custom scan class to hold all relevant scan header information (e.g. scan order, precursor m/z, parent scan information, scan description, polarity) as well as all peak data. Peak data includes ion intensity and m/z and can be extended to include information pertaining to baseline intensity and peak noise if available.

### Monoisotopic Peak Correction

For each MS2 spectra, Monocle determined the parent scan and isolation m/z, then determined the flanking MS^1^ scans for this parent scan (Figure 1A). Peaks within a mass specific window around the isolate precursor were then tracked in each flanking MS1 spectra to encompass the observed precursor envelope (E_O_) as well as an extended window for potential isotopic shifts (up to 14 peaks). E_O_ was determined for either the instrument assigned charge state (default) or individually for each of a user-assigned charge range. A theoretical isotopic envelope (E_T_, up to 7 peaks) was then determined based on the precursor mass using an averagine estimation of the carbon content at a given mass (averagine m/z = 111; averagine carbon count = 5.1)^15^. The relative intensities for E_T_ peaks were estimated using the binomial probability mass function accounting for the relative probability of ^13^C (∼1.1%). E_O_ and E_T_ were scaled to a maximum intensity of 1 and E_O_ was weighted based on the presence of the envelope peaks in the flanking MS1 scans. Monocle then scanned the window surrounding E_O_ for peaks that matched the distribution of E_T_ (Figure 1). The largest dot product of E_O_ and E_T_ was used to determine which observed MS1 peaks belonged to the isotopic window for the precursor. The resulting estimation of the true E_O_ was then used to determine the most likely monoisotopic peak. Finally, Monocle calculated the final monoisotopic m/z based on an intensity weighted average of all flanking MS1 scans.

**Figure 1.**
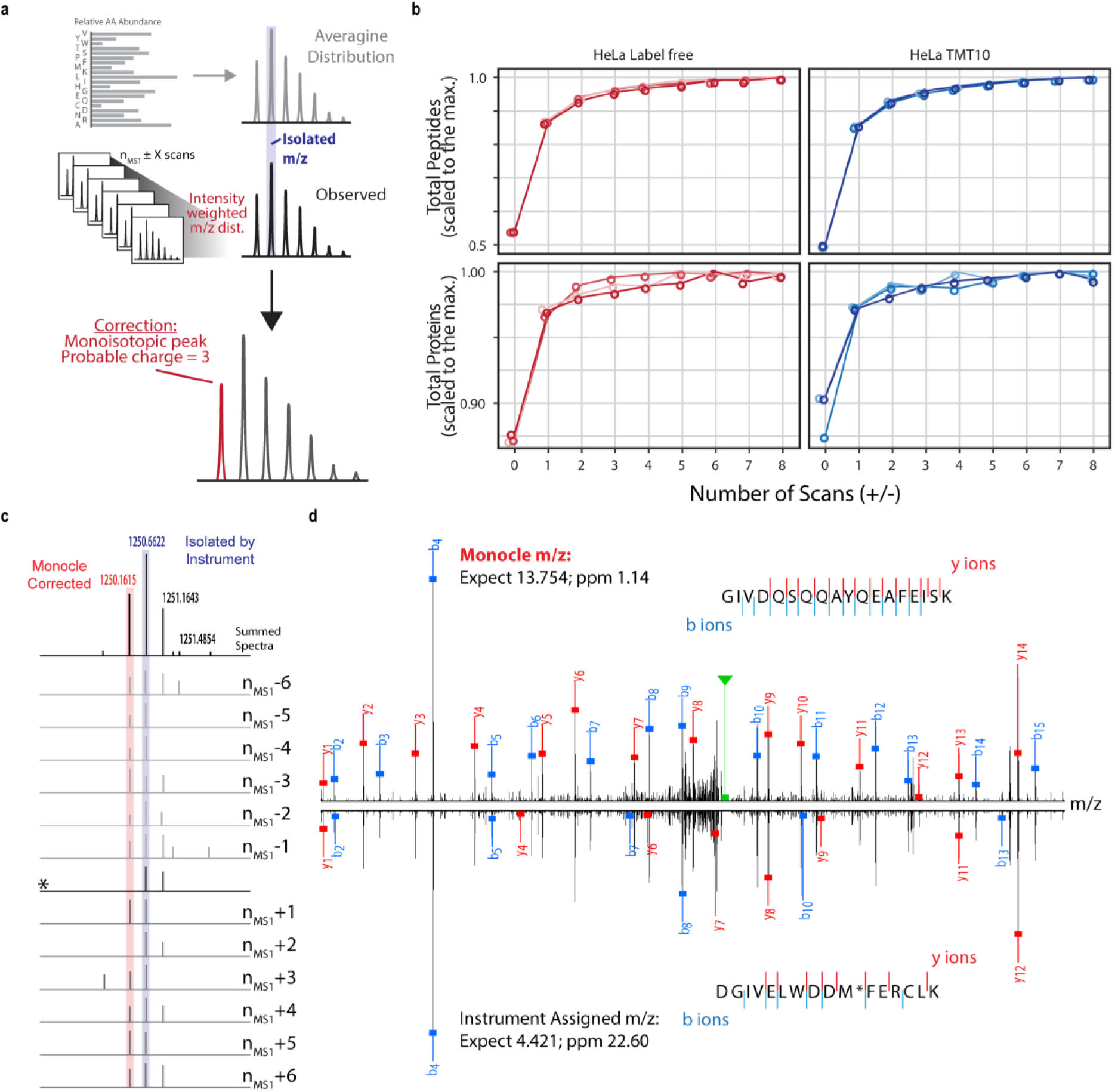
Monocle increases total peptide and protein identifications. A. Workflow overview of the Monocle algorithm. Theoretical isotopic envelopes (E_t_) were estimated based on an averagine distribution. (E_t_) was compared to the observed isotopic envelope (E_o_) for the precursor scan and flanking scans (user defined window). The weighted average of theoretical monoisotopic peak for the best match between E_t_ and E_o_ was then used as the monoisotopic peak. B. Comparison of multiple different window sizes were used to optimize the number of MS1 scans before and after the precursor MS1 scan. At +/- 6 MS1 scans, both peptide and protein curves reached near maximum or maximum sensitivity, respectively. C. Example of combining data from +/- 6 MS1 scans for a given precursor (TMT labeled HeLa sample) to generate a better estimation of the isotopic envelope compared to the precursor MS1 scan (*). D. The resulting PSMs for precursors with correction (top) or without correction (bottom) for the precursor in C.

### Monocle Data Export and Output File Formats

Monocle has been adapted to export mzXML, mzML, and custom csv file formats for implementation with common informatic pipelines^7, 8^. The resulting files are readily searchable through common algorithms (e.g. Sequest^16^, Comet^10^, MaxQuant^17^).

### Comparison to RawConverter

RawConverter (1.1.023) was used with default parameters for data-dependent analysis. The mzXML results were directly used for searching and compared to uncorrected or Monocle corrected data for the exact same raw data files.

### Algorithm Speed Test

Algorithms were tested on a desktop computer running Windows 10 (x64) with an AMD FX-4100 Quad-Core processor (3.60GHz) and 16GB of memory. Processing time included file and scan loading time.

## Results and Discussion

Monocle enabled cross-platform correction of monoisotopic peaks from raw data input. The algorithm and underlying software can be built in either Linux or Windows environments and employed at server scale (CLI) or for individuals (GUI). With either implementation, Monocle read through scans to identify parent and child scans. Precursor masses for the child scan were then used to build estimations of the observed isotopic envelope (E_O_). Monocle identified precursor scans at user defined intervals before and/or after the parent scan (defaulted to +/- parent scans, Figure 1A). These envelopes were compared to a theoretical isotopic envelope (E_T_) based on poly-averagine to approximate ^13^C incorporation. A charge state range could be applied at this stage (e.g. for low resolution MS1 data) and the charge state envelope with the highest dot product when compared to E_T_ was retained. Finally, the monoisotopic peak was calculated based on an intensity-weighted average of the m/z from the parent scan and surrounding scans.

To determine the optimal number of MS1 scans to include for averaging, we compared the total peptide spectral matches (PSMs) and total protein identifications from a human cell line analysis of both label-free and TMT-labeled peptides (Figure 1B). We observed that for both the labeled and unlabeled samples, total PSMs rose quickly to greater 90% of maximal values by incorporating +/- 3 scans and reached near maximum values at +/-6 scans. Furthermore, total protein identifications reached their maximum at +/-6 scans. Based on these data, we proceeded to use +/- 6 MS1 scans (a total of 13 MS1 scans) throughout this work. Using +/- 6 MS1 scans, we observed a nearly 2-fold increase in either label-free or TMT labeled human peptide identifications compared to using no flanking MS1 scans (Figure 1B). Using the +/-6 scan strategy, the mean MS2 processing speed for Monocle was 476.5 +/- 61 Hz (Figure S1, Table S2).

Incorporation of flanking MS1 scans helped to improve monoisotopic peak determination, particularly when misassigned isotopes (Figure 1C). As an example, a TMT-labelled peptide precursor was originally misassigned to a monoisotopic peak of 1250.6622 m/z at z = 2 which matched with a low PSM score and high ppm error to the peptide DGIVELWDDM[ox]FERCLK (Figure 1C, Figure 1D). Averaging of the m/z region for the parent MS1 scan and the twelve flanking MS1 scans, however, makes clear that the monoisotopic peak should have been assigned to 1250.1615 m/z, which Monocle corrected (Figure 1C). Monocle’s monoisotopic peak correction generated a better match (higher expect score and lower ppm error) compared to the original monoisotopic peak assignment, resulting in a PSM to GIVDQSQQAYQEAFEISK (Figure 1D).

We compared the delta mass values for the TMT labeled human peptides and generally observed discrete shifts corresponding to misassignment originally to the wrong precursor isotope (Figure 1B, Figure 2A). Closer inspection revealed tailing around these discrete isotopic shifts (Figure 2B, Figure 2C). Monocle adjustment corrects peak assignments based on the combination of (1) the incorporation of single isotope additions and (2) small, ppm-level shifts (Figure 2A, Figure 2B). First, in keeping with these data, isotopic peak correction has been shown previously to improve the assignment of PSMs for Thermo instruments^2, 3, 5^. Second, we observed that the weighted averaging of peaks from multiple MS1 scans resulted in ppm level shifts (Figure 2B, Figure 2C). These small shifts resulted in more precise ppm error estimates for PSMs using Monocle (Monocle: σ = 0.67; Original: σ = 0.87).

**Figure 2.**
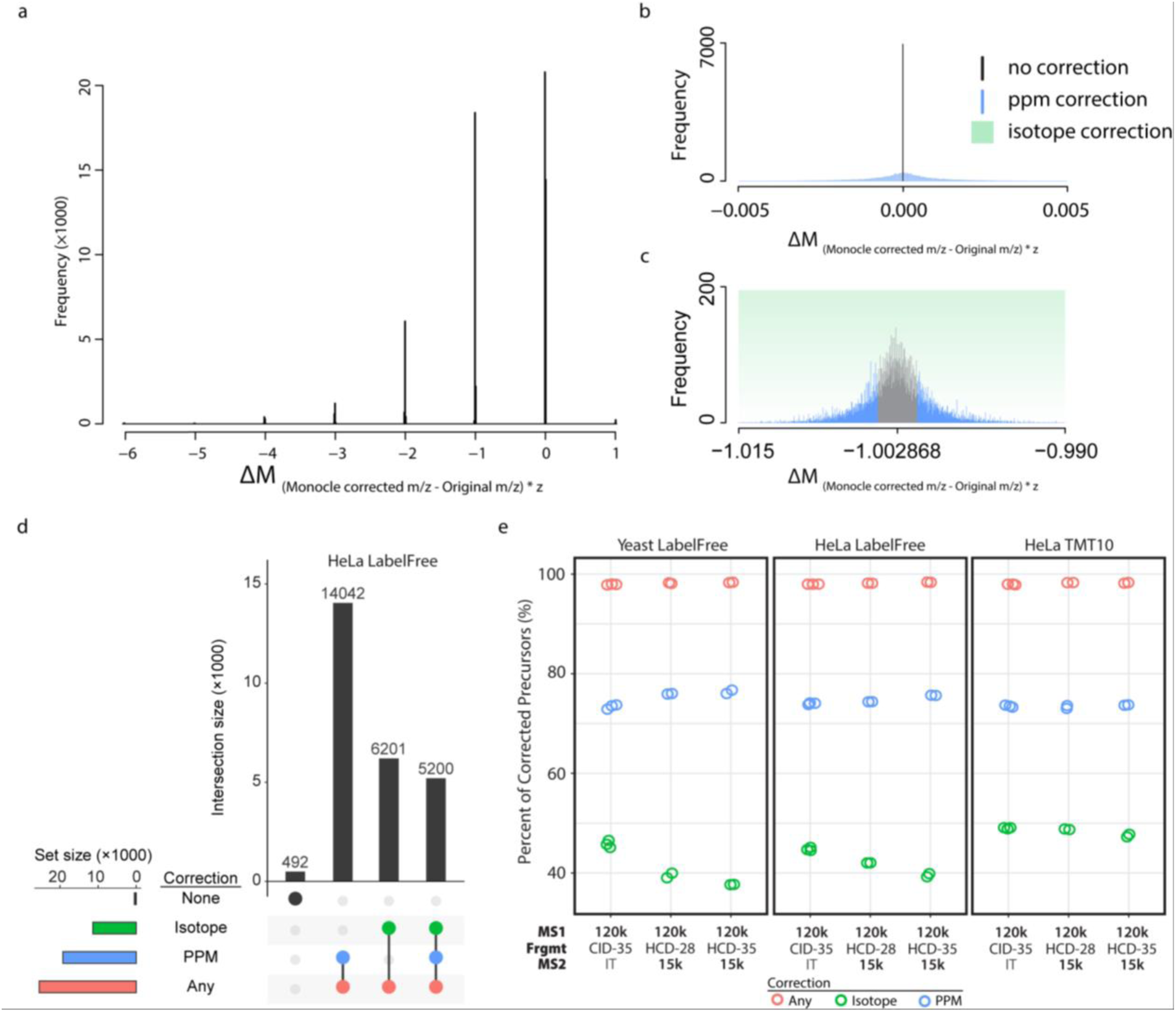
A. Histogram of the delta mass (ΔM) between the Monocle corrected mass and the original, instrument assigned mass based on all unfiltered PSMs. Subsets of the histogram in A at regions around ΔM = 0Da (B) and ΔM = -1Da (C). The histogram in C was centered on the averagine mass defect. D. Comparison of uncorrected precursors to Monocle corrected precursors for all valid peptides revealed two general classes (small ppm deviations and larger isotopic deviations) were highlighted. E Percentage of precursors for valid PSMs corrected by either ppm deviation, isotope deviation or both.

To ensure Monocle was effective for a variety of common instrument settings and applications, it was tested on runs with both high and low-resolution MS/MS workflows. On average 74.5% of precursors were corrected for smaller ppm level shifts, 43.5% of precursors were corrected for isotopic shifts, and 98.3% of precursors were corrected by one or both (Figure 2D, Figure 2E, Figure S2). Differences in types of precursor correction could be explained, in part, by variations in the depth of precursor sampling. Low-resolution workflows sampled lower-abundance precursors which are more likely to be assigned incorrect m/z by instrument software. Interestingly, the maximum intensity across the chromatographic run of peptide precursors significantly affected whether peaks were corrected (p < 0.0001, linear regression; Figure S3). As expected, we also observed a charge state dependence for each correction type (Figure S4). For example, PSMs with 2+ precursors were predominantly corrected with small ppm-level shifts. As charge increased, isotopic correction became a larger proportion of the total corrected precursors (Figure S4). This could partially be explained by lower relative intensities of the ^12^C peaks for larger peptides and high charge state peaks. Small differences in the percentage of ppm shifts between IT-MS2 and FT-MS2 analyses correlated with ppm drift due to instrument performance (IT-MS2: mean ppm error = 0.71; FT-MS2-HCD28: mean ppm error = 0.58).

For a yeast label-free analyzed by the IT-MS2 method, Monocle increased the total number of valid PSMs by 11617 ± 184.18 (14336 ± 424.83 to 25953 ± 419.86) compared to the same analysis with the original, instrument-assigned m/z (Figure 3A, Figure S5A). We explored whether the improvement was due to the relatively low complexity of the yeast proteome by analyzing label-free human peptides derived from HeLa cells, and TMT labeled human peptides. Monocle correction of precursor m/z values for human whole cell lysate resulted in an 82.3% gain in total PSMs for the label-free sample and a 97.7% gain for TMT labelled peptides (Figure 3A). Next, we compared Monocle results to RawConverter results for the exact same underlying spectra. RawConverter was shown to outperform previous implementations of monoisotopic peak correction algorithms^2^. On average, RawConverter improved the total number of identified PSMs by 55% (Figure 3A, Figure S5A). By comparison Monocle improved the total identified PSMs by 85% (Figure 3A, Figure S5A).

**Figure 3.**
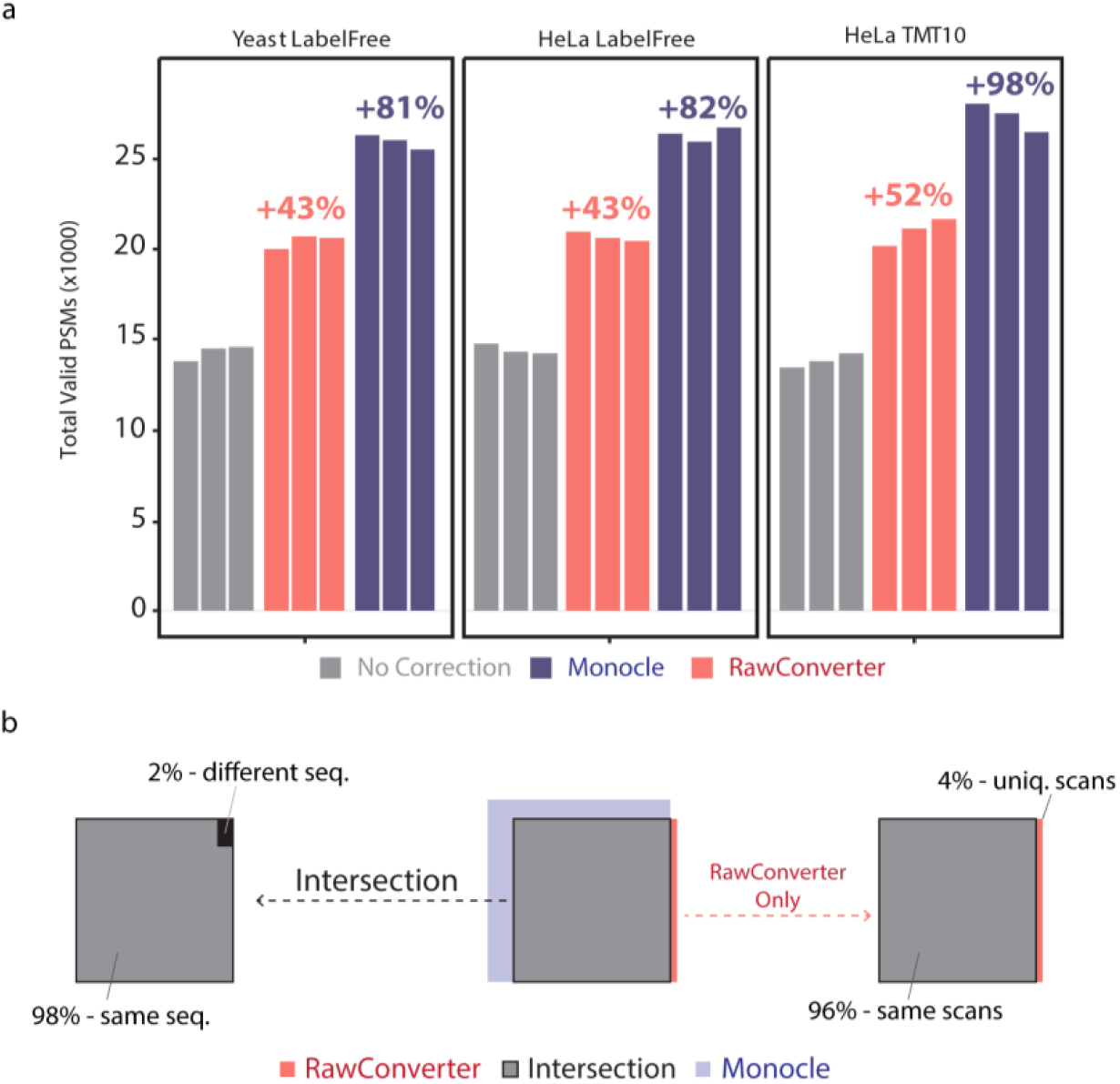
Monoisotopic peak correction increases total PSMs. A, Total valid PSMs for either uncorrected precursors or for monoisotopic peak correction with RawConverter or Monocle. Percentages were the average improvement over uncorrected data. B, Comparison of scans with valid PSMs for Monocle compared to RawConverter. Leftmost: The proportion of the scans that generated a PSM after Monocle and RawConverter that matched to the same peptide sequence. Rightmost: The proportion of RawConverter PSM scans that also generated valid PSMs after Monocle.

We investigated if the same scan generated a valid PSM for the original m/z, the RawConverter m/z or the Monocle corrected monoisotopic peaks (Figure 3B). For the original m/z values, 91% of the PSM generating scans were also identified when RawConverter was used versus 96% with Monocle (Figure S5A). For those scans that generated a PSM for both the original and Monocle m/z values, 99% matched to the same peptide sequence compared to 97% for RawConverter (Figure S5B). Monocle generated PSMs for 96% of PSM scans identified with RawConverter as well (Figure S5C). These data highlight that Monocle can capture the vast majority of valid PSMs identified with the original values or RawConverter values, in addition to adding new, valid PSMs (Figure S6).

Monocle’s marked improvements in total PSMs lead us to test if it could be applied to new acquisition strategies employed on Thermo instruments. We therefore tested the same sample (human TMT-labeled peptides) analyzed in triplicate with or without advanced precursor detection (APD)^18^ or high-frequency asymmetric waveform ion mobility (FAIMS)^19^. Both techniques have been shown to increase the number of viable precursors in an analytical run. APD’s improved on-the-fly monoisotopic peak determination dramatically increased the number of sampled precursors and thereby increase the number of MS2 scans acquired (Figure 4, Figure S7). When we applied Monocle to data generated with APD acquisitions, we observed a similar profile of precursor peak correction as when APD was not applied (Figure 4A). The use of APD increased the total valid PSMs observed (Figure 4B), but APD acquisition was inefficient compared to the canonical acquisition (Figure 4C).

**Figure 4.**
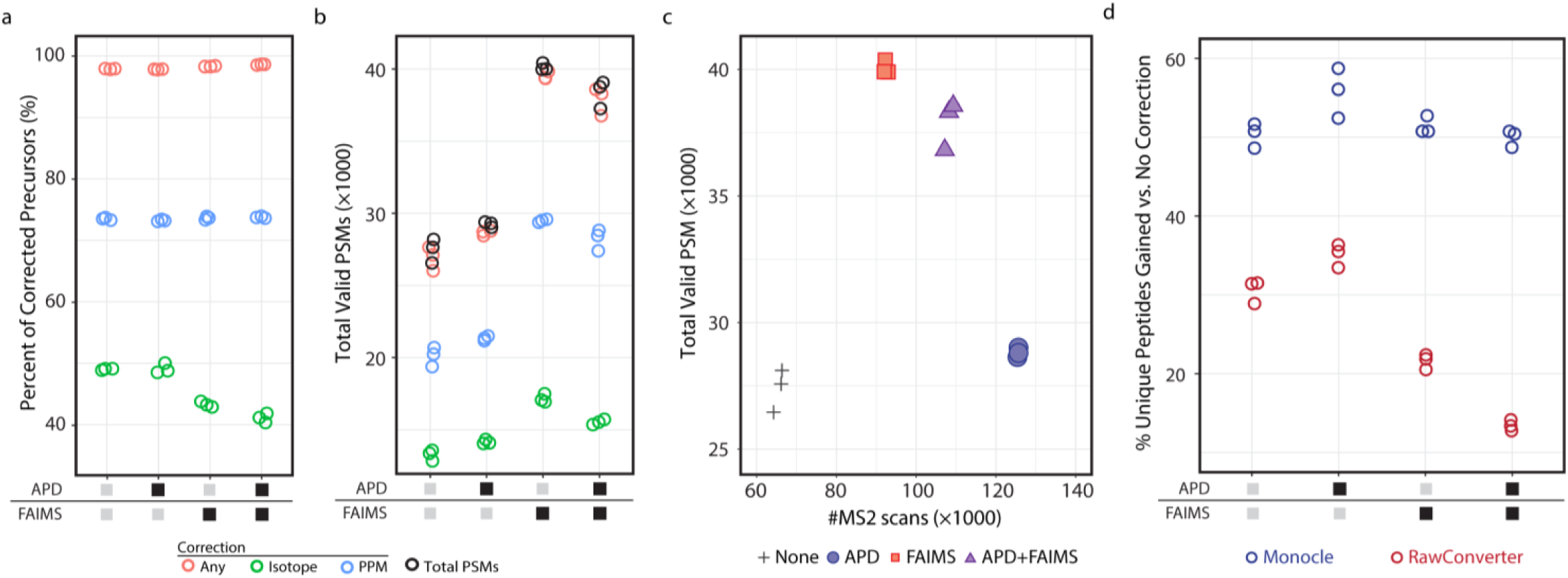
Percentage of precursors corrected based on three different acquisition strategies: APD, FAIMS or FAIMS with APD. B. Total number of valid PSMs identified for each acquisition strategy classified by how Monocle corrected the precursors. C. Success rate of PSMs based on acquisition strategy. D. Comparison of the increased number of unique peptide identifications observed for Monocle and RawConverter based on different acquisition strategies compared to the original m/z value analysis.

The application of FAIMS, which can reduce co-isolation of overlapping precursors^19^, reduced the percentage of isotope corrected precursors by 5.5% without APD and 8.1% with APD (Figure 4A). Moreover, FAIMS increased the number of valid PSMs for the same sample and injection amount (Figure 4B) and provided more efficient conversion of acquired MS2s to valid PSMs (Figure 4C). When combined, FAIMS and APD identified more valid PSMs compared to either canonical acquisition or APD-alone and provided intermediate efficiency (Figure 4C).

Both Monocle and RawConverter robustly improved the total identified PSMs for all four acquisition strategies (Figure 4D, Figure S8). Monocle outperformed RawConverter for the canonical acquisition and methods that incorporated FAIMS (Figure 4D). Monocle and RawConverter generated similar total number of PSMs when APD alone was used (Figure S6), however Monocle generated more unique peptide identifications (Figure 4D).

In this work we show the utility of monoisotopic peak estimation and correction to improve the sensitivity of proteomics identification. The platform presented here, Monocle, has been released as a cross platform, open-source library to enable its use in the community. The command line and graphical interfaces will enable adoption in a wide variety of pipelines moving forward. Finally, we showed that Monocle can nearly double the total peptide spectral matches for label-free and TMT-labeled proteomes and that this improvement extends to modern acquisition strategies, e.g. APD and FAIMS.

## Supporting Information

### Tables

Table S1 – Comparison of acquisition methods used in this study.

Table S2 – Precursor processing time.

### Figures

Figure S1 – Monocle raw file processing speeds.

Figure S2 – Precursor correction across species and labeling methods.

Figure S3 – Effect of maximum precursor intensity (chromatographic apex) on isotopic peak correction for TMT labeled human peptides.

Figure S4 – Classification of precursor correction and the effect of charge on correction class.

Figure S5 – The percentage of precursors matches when comparing monoisotopic peaks estimation methods across species and derivatization techniques.

Figure S6 – Relative improvements in PSMs using Monocle or RawConverter compared to No Correction.

Figure S7 – Comparison of unfiltered APD and FAIMS correction data.

Figure S8 – The percentage of precursors matches when comparing monoisotopic peaks estimation methods based on data acquired with APD, FAIMS, or APD and FAIMS.

## Acknowledgements

The authors would like to thank past and present members of the Gygi lab for constructive critiques and technical advice. Specifically, the authors would like to acknowledge work by Drs Josh Elias and Sean Beausoleil on earlier iterations of monoisotopic peak correction. We acknowledge the following sources of funding: GM067945 to S.P.G.

