## Supporting Information for "Improved Monoisotopic Mass Estimation for Deeper Proteome Coverage"

**Contents**

*Tables*
Table S1 – Comparison of acquisition methods used in this study.
Table S2 – Precursor processing time.

*Figures*
Figure S1 – Monocle raw file processing speeds.
Figure S2 – Precursor correction across species and labeling methods.
Figure S3 – Effect of maximum precursor intensity (chromatographic apex) on isotopic peak correction for TMT labeled human peptides.
Figure S4 – Classification of precursor correction and the effect of charge on correction class.
Figure S5 – The percentage of precursors matches when comparing monoisotopic peaks estimation methods across species and derivatization techniques.
Figure S6 – Relative improvements in PSMs using Monocle or RawConverter compared to No Correction.
Figure S7 – Comparison of unfiltered APD and FAIMS correction data.
Figure S8 – The percentage of precursors matches when comparing monoisotopic peaks estimation methods based on data acquired with APD, FAIMS, or APD and FAIMS.

**Table S1** Comparison of acquisition methods used in this study.

| Method Name | MS1 Analyzer | MS1 Resolution | MS2 Analyzer | MS2 Resolution | Fragmentation Type | Fragmentation Energy | FAIMS DV | FAMS CV |
|---|---|---|---|---|---|---|---|---|
| (*Default*) | Orbitrap | 120,000 | Ion trap | Rapid | CID | 35% | - | - |
| **Low Res** | Orbitrap | 120,000 | Ion trap | Rapid | CID | 35% | - | - |
| **High-Res1** | Orbitrap | 120,000 | **Orbitrap** | **15,000** | **HCD** | **28%** | - | - |
| **High-Res2** | Orbitrap | 120,000 | **Orbitrap** | **15,000** | **HCD** | 35% | - | - |
| **APD** | Orbitrap | 120,000 | Ion trap | Rapid | CID | 35% | - | - |
| **FAIMS** | Orbitrap | 120,000 | Ion trap | Rapid | CID | 35% | **-5000V** | **-40V**<br>**-60V**<br>**-80V** |
| **FAIMS & APD** | Orbitrap | 120,000 | Ion trap | Rapid | CID | 35% | **-5000V** | **-40V**<br>**-60V**<br>**-80V** |

**Table S2** Precursor processing time. LF – Label-free. Methods described in Table S1.

| Label | Species | Method | Rep. | Total Time (min) | Total Time (sec) | MS2 Count | Monocle Speed (MS2/sec) |
|---|---|---|---|---|---|---|---|
| **LF** | Yeast | Low Res | 1 | 2.35 | 141 | 62754 | 445 |
| **LF** | Yeast | Low Res | 2 | 2.37 | 142 | 62231 | 438 |
| **LF** | Yeast | Low Res | 3 | 2.32 | 139 | 60602 | 436 |
| **LF** | Yeast | High-Res1 | 1 | 1.32 | 79 | 43510 | 551 |
| **LF** | Yeast | High-Res1 | 2 | 1.32 | 79 | 44027 | 557 |
| **LF** | Yeast | High-Res2 | 1 | 1.27 | 76 | 43597 | 574 |
| **LF** | Yeast | High-Res2 | 2 | 1.32 | 79 | 43504 | 551 |
| **LF** | Human | Low Res | 1 | 3.13 | 188 | 71934 | 382 |
| **LF** | Human | Low Res | 2 | 3.10 | 186 | 72375 | 389 |
| **LF** | Human | Low Res | 3 | 3.28 | 197 | 72404 | 368 |
| **LF** | Human | High-Res1 | 1 | 1.48 | 89 | 45998 | 517 |
| **LF** | Human | High-Res1 | 2 | 1.48 | 89 | 46226 | 519 |
| **LF** | Human | High-Res2 | 1 | 1.48 | 89 | 45663 | 513 |
| **LF** | Human | High-Res2 | 2 | 1.57 | 94 | 45803 | 487 |
| **TMT10** | Human | Low Res | 1 | 3.00 | 180 | 68317 | 380 |
| **TMT10** | Human | Low Res | 2 | 3.05 | 183 | 68022 | 372 |
| **TMT10** | Human | Low Res | 3 | 3.00 | 180 | 66070 | 367 |
| **TMT10** | Human | High-Res1 | 1 | 1.52 | 91 | 45021 | 495 |
| **TMT10** | Human | High-Res1 | 2 | 1.53 | 92 | 45058 | 490 |
| **TMT10** | Human | High-Res2 | 1 | 1.52 | 91 | 44763 | 492 |
| **TMT10** | Human | High-Res2 | 2 | 1.58 | 95 | 44677 | 470 |
| **TMT10** | Human | APD | 1 | 4.73 | 284 | 135957 | 479 |
| **TMT10** | Human | APD | 2 | 4.78 | 287 | 136137 | 474 |
| **TMT10** | Human | APD | 3 | 4.87 | 292 | 136086 | 466 |
| **TMT10** | Human | FAIMS | 1 | 3.03 | 182 | 93741 | 515 |
| **TMT10** | Human | FAIMS | 2 | 3.18 | 191 | 93838 | 491 |
| **TMT10** | Human | FAIMS | 3 | 3.20 | 192 | 93976 | 490 |
| **TMT10** | Human | APD+FAIMS | 1 | 3.68 | 221 | 114436 | 518 |
| **TMT10** | Human | APD+FAIMS | 2 | 3.55 | 213 | 114511 | 538 |
| **TMT10** | Human | APD+FAIMS | 3 | 3.60 | 216 | 114877 | 532 |

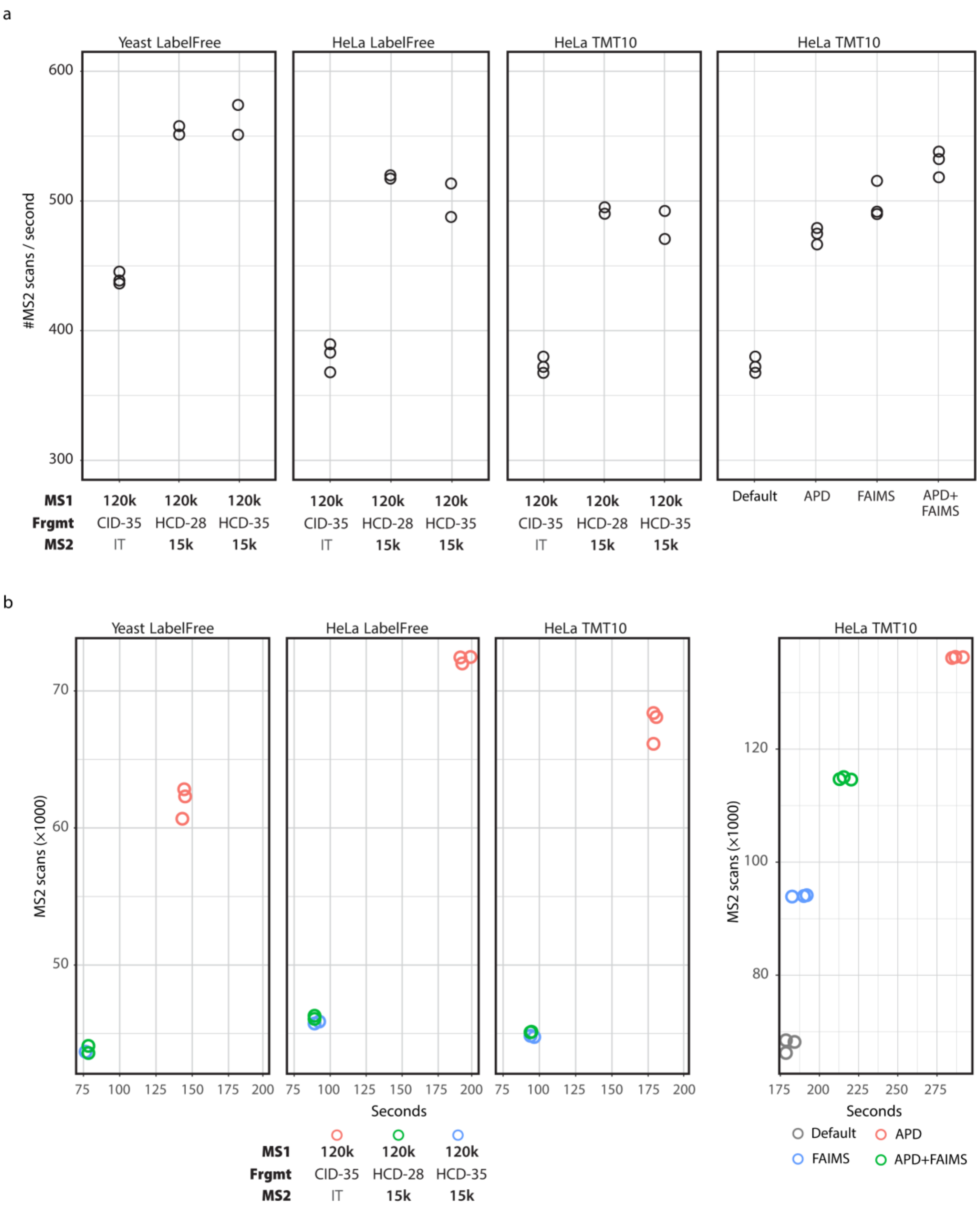


**Figure S1** Monocle raw file processing speeds. (A) Processing speed for each method and replicate across methods in Table S1. (B) Total processing time (seconds) plotted against the total number of scans acquired for each sample analysis.

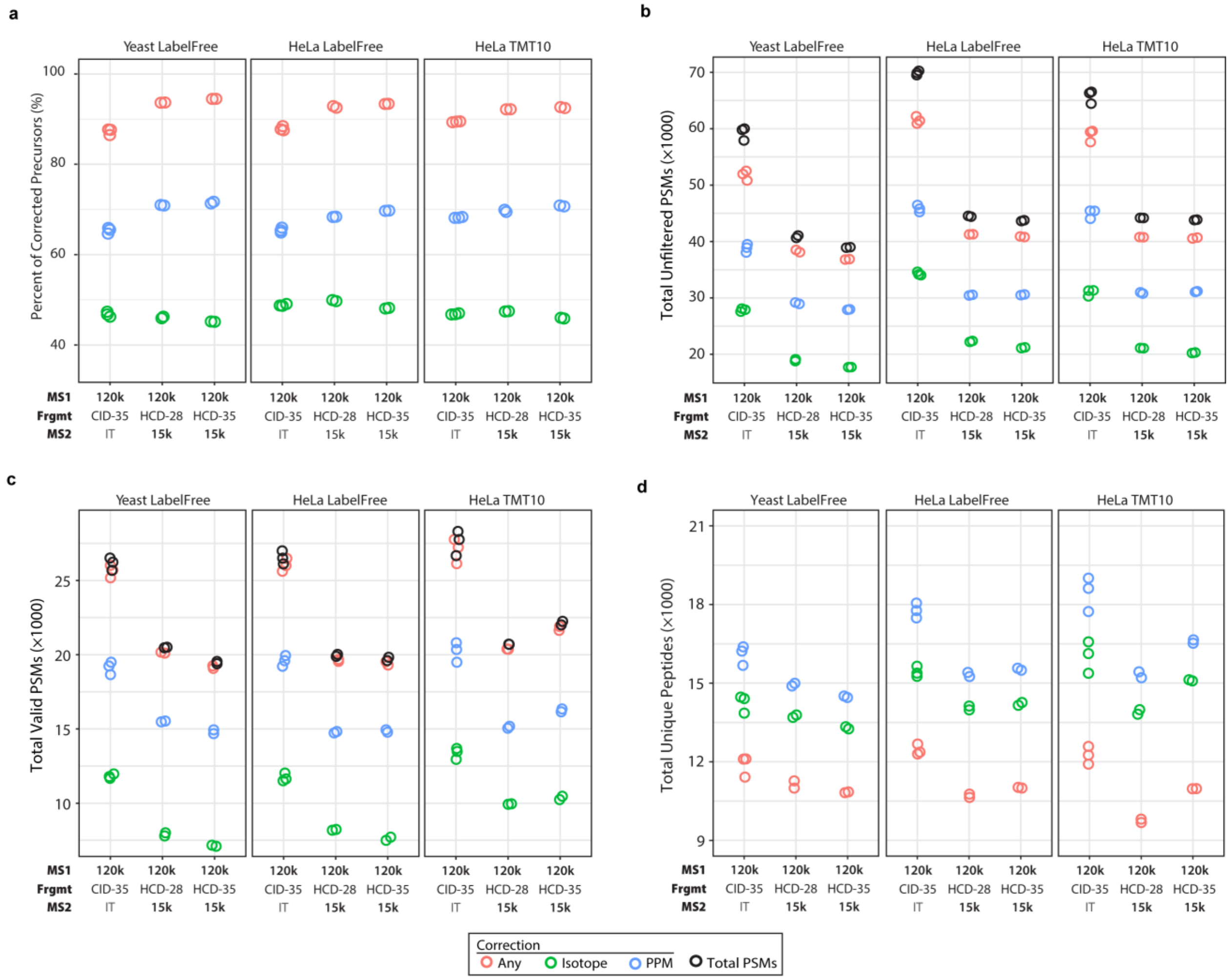


**Figure S2** Precursor correction across species and labeling methods. (A) Percentage of corrected precursors for each method type and species. (B) Total number of unfiltered PSMs for which the monoisotopic peak was corrected by each criterion. The total number of PSMs is shown as black open circles. (C) Total number of valid PSMs (less than 1% peptide and protein FDR) for which the monoisotopic peak was corrected by each criterion. The total number of PSMs is shown as black open circles.

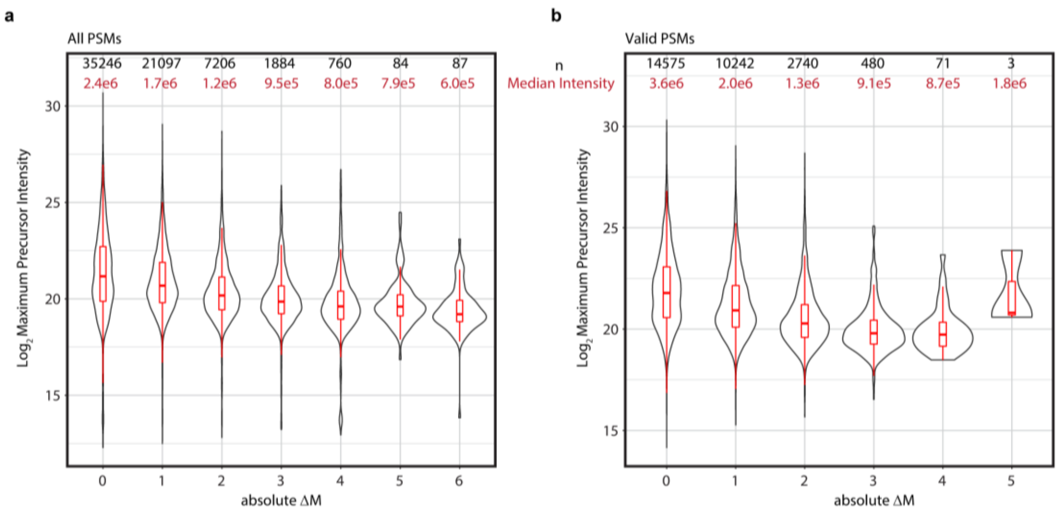


**Figure S3** Effect of maximum precursor intensity (chromatographic apex) on isotopic peak correction for TMT labeled human peptides. Corrected precursors were binned based on absolute delta mass into either the $^{12}C$ or isotopic bins. A significant negative trend was observed (linear model) indicating an inverse relationship between delta mass and maximum precursor intensity. (A) Unfiltered PSMs; (B) Valid PSMs (< 1% peptide and protein FDR).

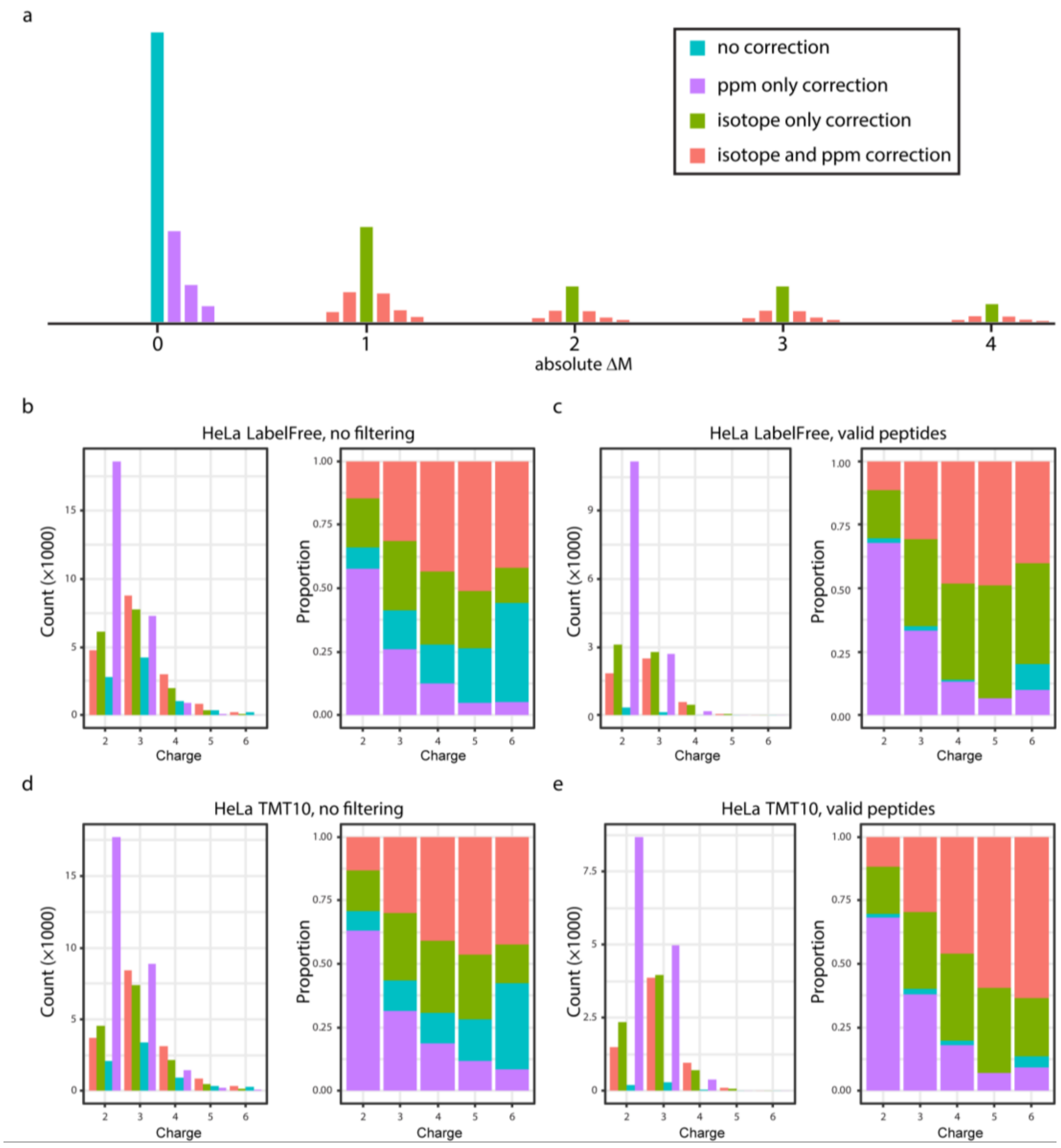


**Figure S4** Classification of precursor correction and the effect of charge on correction class. (A) Diagram of precursor correction classification. (B) Example of precursor correction based on PSM charge showing the total corrected precursors and the relative proportion. (B) and TMT-labeled sample for (C) a human label-free sample without FDR filtering, (D) a human TMT-labeled sample without FDR filtering, and (E) a human TMT-labeled sample with FDR filtering.

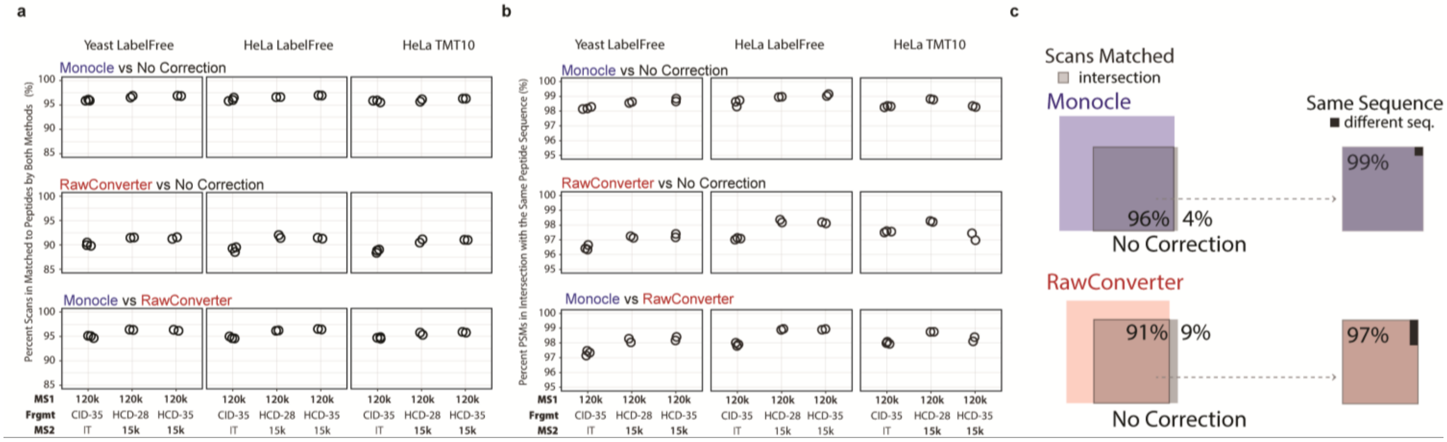


**Figure S5** The percentage of precursors matches when comparing monoisotopic peaks estimation methods across species and derivatization techniques. (A) For scans that generated a PSM with both compared correction methods, the percentage of precursors corrected by Monocle compared to no correction, RawConverter compared to no correction. Percentage and Monocle compared to RawConverter. (B) For scans that generated a PSM with the same sequence, the percentage of precursors corrected by Monocle compared to no correction, RawConverter compared to no correction. Percentage and Monocle compared to RawConverter. (C) Comparison of scans with valid PSMs for Monocle and RawConverter compared to No Correction.

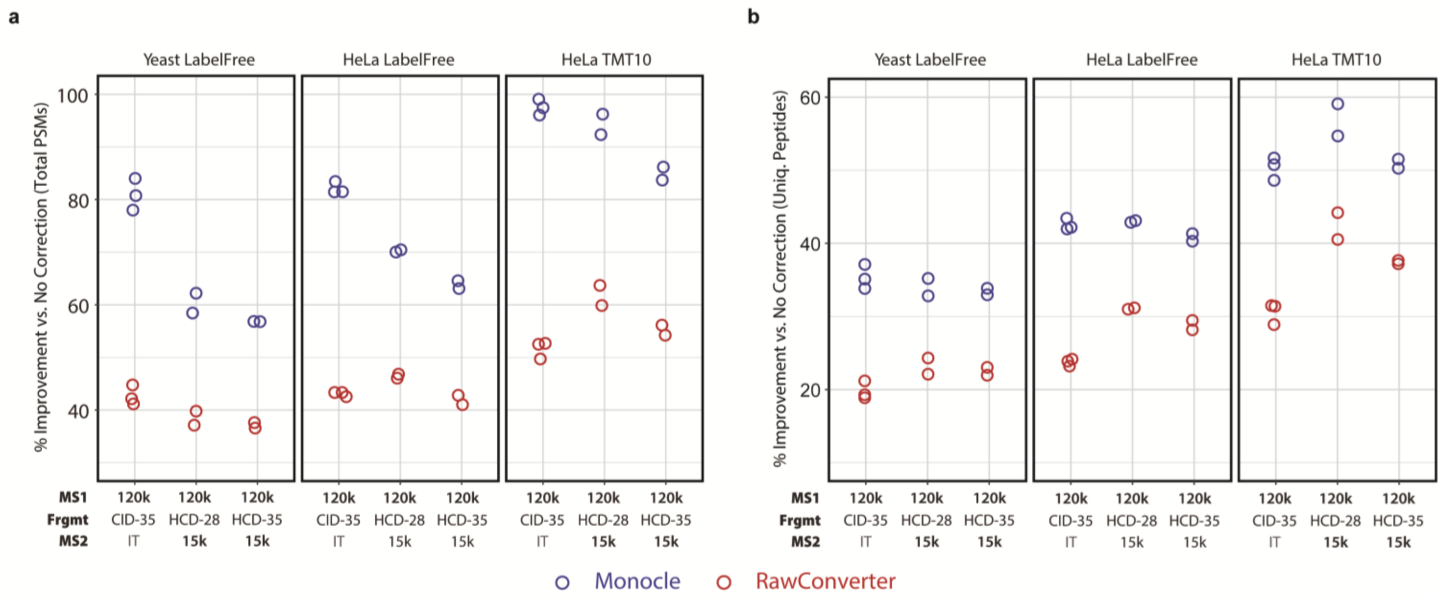


**Figure S6** Relative improvements in PSMs using Monocle or RawConverter compared to No Correction. (A) Total valid PSMs. (B) Unique peptides.

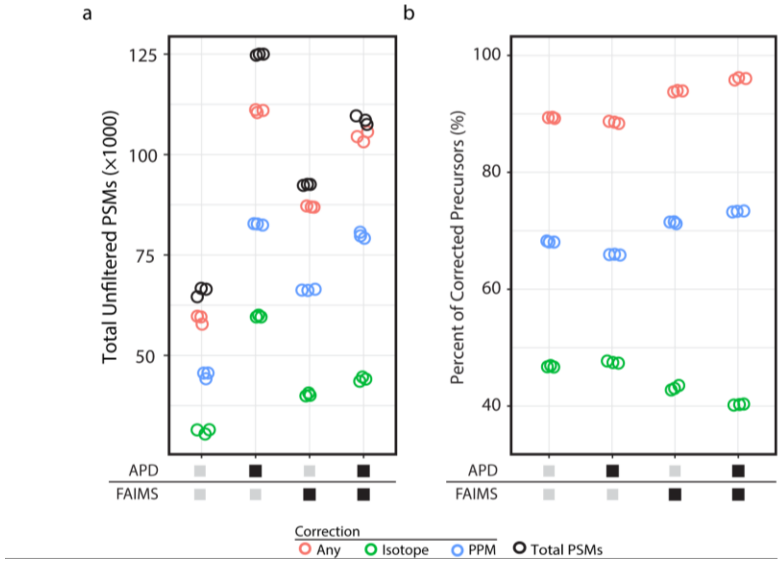


**Figure S7** Comparison of unfiltered APD and FAIMS correction data. (A) Total unfiltered peptide spectral matches (PSMs) for each acquisition method. (B) Percentage of precursors for which the monoisotopic peak estimation was corrected using Monocle. PSM precursor correction types are shown.

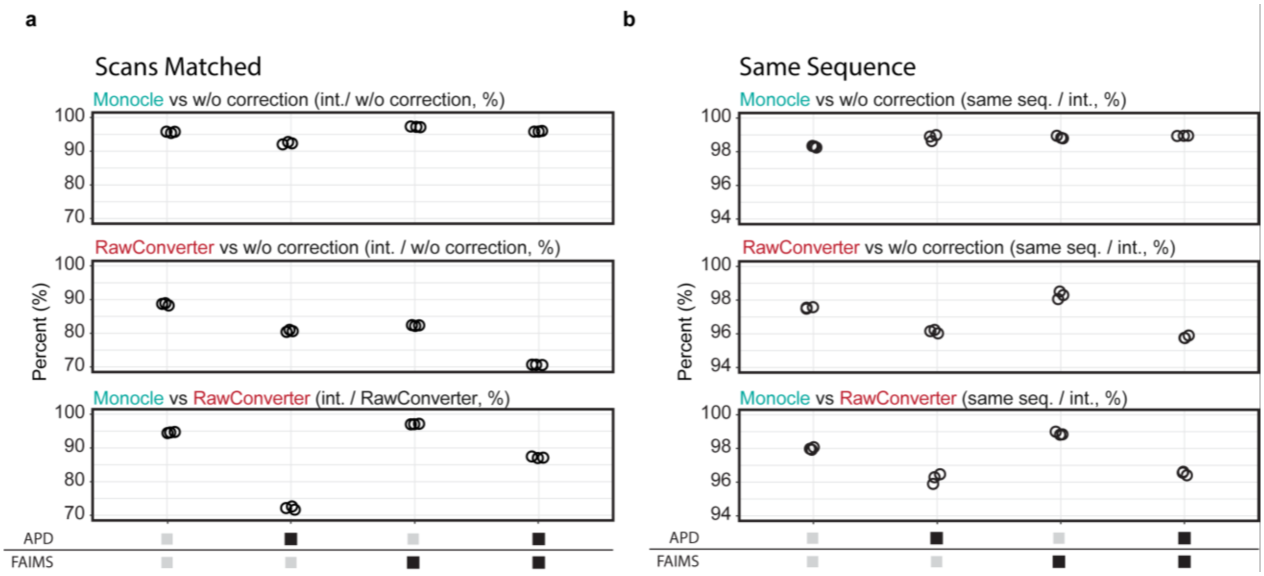


**Figure S8** The percentage of precursors matches when comparing monoisotopic peaks estimation methods based on data acquired with APD, FAIMS, or APD and FAIMS. (A) For scans generated a PSM with both compared correction methods, the percentage of precursors corrected by Monocle compared to no correction, RawConverter compared to no correction. Percentage and Monocle compared to RawConverter. (B) For scans that generated a PSM with the same sequence, the percentage of precursors corrected by Monocle compared to no correction, RawConverter compared to no correction. Percentage and Monocle compared to RawConverter.
